# Is passive dispersal informed? - Experimental evidence for decision-making in phytophagous arthropods

**DOI:** 10.1101/2025.08.30.671941

**Authors:** Kamila Zalewska, Anna Skoracka, Dries Bonte, Ewa Puchalska, Mariusz Lewandowski, Lechosław Kuczyński

## Abstract

Animals must acquire and decode information to make the right decisions. While active dispersers can evaluate habitats *en route*, passive dispersers can only control their departure timing. Although the passive strategy is ubiquitous among arthropods, the mechanisms behind their take-off decisions remain poorly understood. We tested whether host niche breadth shapes passive dispersal in phytophagous mites by exposing them to host-derived kairomones and measuring departure rates. Using experimentally evolved specialist and generalist lineages, we found that dispersal depends more on the context in which cues are encountered that on the kairomones themselves. Host specialisation strongly shaped responses: mites left plants more readily when exposed to unfamiliar hosts, with generalists dispersing over twice as often as specialists. Increased number of unfamiliar kairomones strongly inhibited generalists dispersal but barely affected specialists. This suggests specialists use environmental novelty to trigger exploration, whereas generalists need multiple cues to confirm host suitability, revealing a trade-off between host range and environmental sensitivity.

## INTRODUCTION

Receiving, decoding and using information correctly is a prerequisite for animals to make the right decisions. One of the key decisions, with a significant impact on fitness, is the habitat choice. In an ever-changing environment, animals are forced to track down the optimal habitat and constantly decide whether to stay or disperse. These decisions affect their future interactions with the components of the environment, such as food, shelter, mates, competitors, and predators. Species that move actively can engage in prospecting potential settlement locations and therefore have the opportunity to weigh up the habitat quality of potential new locations against that of their current habitat. This prospecting for personal and social information allows them to make informed decisions about departure, movement and settlement (Clobert et al. 2009; Delgado et al. 2014; McPeek et al. 2024; Ponchon & Travis 2022; Thierry et al. 2024; Usinowicz & O’Connor 2023).

While many species actively decide to depart, they rely on biotic or abiotic vectors, such as water or air currents, to travel to and colonise new locations. Spiders, for instance, engage in passive ballooning, but they actively adopt a body position that facilitates wind or vector take-up (Cho et al. 2018; Montes & Gleiser 2024). From an individual perspective, passive dispersal is extremely risky due to the associated costs during the transfer, such as resource limitations and the depletion of energy reserves during the time spent in an unsuitable landscape. However, these costs may be offset by the massive production of dispersing stages, which allow for bet hedging in uncertain environments. The costs of failing to reach a suitable habitat can indeed be particularly high in a heterogeneous landscape with sparsely distributed habitats. In such a landscape, theory predicts that dispersal would be selected against (Bonte et al. 2012; Fronhofer et al. 2024; Henriques-Silva et al. 2015). On the other hand, the lack of gene flow would increase interspecific competition or inbreeding, both of which are known to favour dispersal again. Importantly, dispersal is usually not a fixed strategy, but rather conditional to local environmental factors such as e.g., kin competition. This can lead to strategies that depend on density, resources or body condition, even when the costs are high (Bonte et al. 2010, 2012; Gros et al. 2006; Poethke & Hovestadt 2002).

To reduce these costs, the information-use can evolve as a fitness-optimising strategy (Mortier et al. 2019), of course pending the costs of receiving and processing, and the reliability of the information received. As passively dispersing organisms cannot engage in prospecting, this information should be received and processed during the departure phase (Bocedi et al. 2012; Delgado et al. 2014; Kiedrowicz et al. 2017).

Overall, dispersal is favoured when the fitness benefits of moving away from one’s place of birth outweigh the costs of remaining there. Therefore, organisms must weigh up the expected fitness benefits of moving away versus staying at home. Individual heterogeneity in these expectations will then induce individual variation in the dispersal strategies (Bonte et al. 2014; Hamilton & May 1977). As population sizes increase, or as resources become limited or the habitat deteriorates, dispersal motivation is therefore expected to increase (Banks et al. 2024; Bonte et al. 2012; Laska et al. 2019).

Crucially, fitness expectations depend directly on the degree to which an organism’s phenotype matches its environment, i.e. its level of local adaptation. Individuals that are strongly adapted to their local environment are known as ecological specialists. When these environmental conditions of local adaptation are sparse, a joint evolution of specialisation and philopatry is theoretically predicted (Ravigne et al. 2024). The basic assumption is that specialists should be poor dispersers to avoid landing in unsuitable habitats, whereas generalists should have high dispersal abilities to take advantage of suitable opportunities for settlement. While some empirical studies have confirmed this expectation (e.g. Freedman et al. 2020; Kneitel 2018; Stevens et al. 2012, 2013), others have not (e.g. Alzate et al. 2017; Dapporto & Dennis 2013; Van Zandt & Mopper 1998). This discrepancy is probably due to the fact that the spatio-temporal dynamics of the conditions to which species have adapted are often overlooked (Ravigne et al. 2024). However, it remains to be confirmed whether the degree of specialisation affects emigration in passive dispersers or if there are differences in the ability of specialists and generalists to perceive information from both the local environment and potential immigration locations. Under optimality, we would expect both specialists and generalists to correctly interpret cues from a familiar environment when making dispersal decisions. Alternatively, since generalists are more likely to land in suitable habitats than specialists, we can hypothesise that generalists do not invest as much in receiving signals as specialists do, as this incurs costs in terms of morphological and physiological adaptations, as well as time spent receiving and recognising cues. If this scenario is true, we should observe specialists responding more strongly to cues than generalists, and overall lower dispersal motivation.

In summary, in order to answer the question of whether departures are informed in passively dispersing organisms, it is crucial to assess whether organisms respond proportionally to the strength of the signal they receive. It is equally crucial to consider the context of the current and future environments. Otherwise, we will not be able to demonstrate whether the information is decodable or selective. Furthermore, examining the extent of ecological specialisation can provide additional insight into whether the interplay between specialisation and dispersal involves an element of information use.

Here, we address above the questions, using passively dispersing herbivorous arthropods belonging to lineages that have been adapted through over a hundred of generations of experimental evolution to different plant species regimes, and thus they differ in their degree of host specialisation. Lineages that evolved in a fluctuating environment, consisting of temporally alternating wheat and barley, increased their ability to explore different plant resources, including species not encountered during experimental evolution.

Consequently, they represent flexible host-use generalist phenotypes. In contrast, a stable one-host environment (wheat) led to wheat-specialised phenotypes with decreased performance on other hosts (Skoracka et al. 2022). In the present study, different plant species, familiar or unfamiliar, have served as either current environments, from which animals may have emigrated, or target environments, which they would reach after dispersal. Specifically, we ask whether the departure rate is influenced by:

i. information from potential target host environments in the context of the current host environment;
ii. the degree of host specialisation;
iii. the signal-to-noise ratio, i.e. the number of cues provided by the target environment.

Generally we showed that the response to information about the target environment is selective, but also highly context dependent. The decision to emigrate depends not only on environmental information, but is highly modulated by the degree of host specialisation.

## MATERIAL AND METHODS

### The study system

We used an obligate phytophagous mite *Aceria tosichella* (the wheat curl mite, hereafter WCM), which is an economically important pest of agricultural crops (Box 1). The WCM represents a cryptic species complex consisting of several divergent genotypes. We performed our experiment on the MT-1 genotype, which was identified by DNA barcoding (Skoracka et al. 2018a). The WCM MT-1 is one of the most serious pests of wheat worldwide, causing significant losses through direct feeding and by transmitting plant viruses, which increases its invasive potential and colonisation success. Its short development time (six to seven days at 27°C), rapid multiplication capacity, high dispersal ability, and thermal tolerance make it a suitable organism for laboratory manipulation and amenable to experimental evolution (Karpicka-Ignatowska et al. 2021; Kuczyński et al. 2016; Laska et al. 2021; Skoracka et al. 2018b, 2022). The WCM mites are dispersed passively by wind currents. When exposed to wind, they respond quickly, indicating either the propensity of dispersal or a lack of it (Kuczyński et al. 2020; Laska et al. 2019).

### Experimental evolution

We used lineages of WCM MT-1 that differed in their host specialisation. To avoid background noise caused by genetic heterogeneity among these lineages, we leveraged rigorous, replicated experimental evolution to obtain lineages that were sufficiently different yet comparable with respect to the trait of interest (i.e. ecological specialisation). These lineages were obtained through over 100 generations of experimental adaptation to different host regimes, starting from the same baseline colony.

The baseline colony was established in November 2017 using 26 WCM MT-1 populations collected from wheat fields in nine geographically distant locations across Poland. To confirm their MT-1 genotype, individuals from all of the field-collected populations were barcoded using mtDNA COI. Approximately 1,000 adult females from each population were then combined to form the baseline colony. This colony was then maintained under constant conditions (22–24°C, 12:12 D/N, 40% RH) for four weeks, after which the individuals were used in the experimental evolution. This procedure ensured the genetic diversity within the baseline colony enabling generalists and specialists to be selected from the same gene pool.

In order to investigate the effects of different host-use regimes, two evolutionary treatments were established: a specialist regime, in which populations were maintained exclusively on wheat (*Triticum aestivum*); and a generalist regime, in which populations alternated between wheat and barley (*Hordeum vulgare*). Six independent evolving lineages (replicates) were initiated for each regime. Each lineage began with 300 WCM MT-1 individuals, which were sampled from a baseline colony and transferred to clean potted wheat plants.

All lineages were maintained under controlled conditions (27°C, 16:8-hour light:dark cycle, 60% relative humidity) in separate isolators to prevent cross-contamination. Every three weeks—corresponding to approximately three generations at 27°C (Karpicka-Ignatowska et al. 2021)—around 300 individuals from each lineage were transferred to fresh plants according to their assigned regime: wheat-only for specialists, and alternating wheat/barley for generalists.

One of the specialist lineages failed to survive during the experiment. Once evolution had run its course, the fitness of five specialists and six generalists was assessed across a variety of plant species in order to verify their host specialisation. The alternating host environment resulted in an increased host niche breadth, encompassing plant species that had not been encountered during experimental evolution. By contrast, the constant host environment led to specialised wheat phenotypes that performed poorly on barley. All lineages within each evolutionary regime exhibited a consistent pattern of host adaptation (Skoracka et al. 2022).

### Experimental design

We designed an experimental setup consisting of:

i. the current environment, i.e. the plant species on which the mites were placed and remained throughout the experiment; and
ii. the target environment, i.e. the plant species, with which the mites did not come into the direct contact, but from which the olfactory cues (kairomones) came. We only used kairomones emitted by the plants, with no mechanical interference or induction by feeding mites. This was done deliberately to avoid influencing factors such as competition or population density, and to provide a pure signal of the presence of a particular plant species.

Both, the current and target environments were either:

a. familiar, i.e. an environment composed of plant species or cues derived from them, with which the mites had contact during experimental evolution, or
b. unfamiliar, i.e. an environment composed of plant species or cues derived from them, with which the mites had no contact during their evolution.

To assess whether a departure is informed, it is necessary to provide information about the current and potential target environments, and to measure the response – that is, the organisms’ willingness to be taken off. To distinguish whether a mite’s willingness to depart is due to its greater overall ability to disperse or to cue recognition ability, it is essential to compare responses in control groups without cues and experimental groups with cues. If the responses of mites are similar in the control and experimental groups, this suggests that they had not evolved informed dispersal; however mites generally have high dispersal potential.

To achieve this, we exposed mites to wind currents carrying different kairomones using an olfactometer. The following plant species were used to create the current and target environments: wheat (W), barley (B), smooth brome, *Bromus inermis* (S), and oat, *Avena sativa* (O). Specialist and generalist individuals (from five and six lineages, respectively) were exposed to cues from the current and target environments, and their response was measured as the proportion of individuals that dispersed. Specifically, variants containing wheat (W) provided a familiar environment for specialists, and variants containing both wheat (W) and barley (B) provided a familiar environment for generalists. Other plant species provided an unfamiliar environment.

Specifically, we conducted the following experiments:

i. Experiment 1: Dispersal in response to a single kairomone in the context of the current environment and host specialisation. This was achieved by recording the departure rates of specialists and generalists when they settled on W, B, or S, and were exposed to kairomones produced by W, B, O, or S, or clean air (the control group).
ii. Experiment 2: Dispersal in response to signal noise through a mixture of kairomones from the target environment. This was achieved by recording the departure rates of specialists and generalists as they settled on W and were exposed to mixtures containing two or three kairomones: W-S, S-O, B-S-O, or W-S-O. The mixtures contained either only unfamiliar cues or a combination of familiar and unfamiliar cues.

Each experiment was performed separately for the specialist and generalist lineages. See Supporting Information 1, Table S1 for details about experimental variants.

The olfactometer was made up of tightly connected elements that allowed the kairomones to reach the experimental arena, where the mites’ reactions were tested. These elements included: an air pump that acted as kairomones generator, and a chamber containing a plant that served as the source of the kairomones. There were also chambers filled with activated charcoal to provide clean air input or output (Figs S1-S2 in Supporting Information 1).

The experimental arena consisted of a 1 cm^2^ plant fragment placed on agar blocks prepared from a modified *in vitro* culture medium (Karpicka-Igantowska et al. 2019; Murashige and Skoog 1962), which represented the current environment. In Experiment 1, the current environment consisted of wheat (a familiar to both specialists and generalists) or barley (a familiar to generalists only) or brome (an unfamiliar to both specialists and generalists). In Experiment 2, the current environment contained only wheat. At least ten WCM MT-1 females were transferred to the arena and acclimatised for 30 minutes. The arena was then placed in the olfactometer connected to an Olympus SZX7 stereomicroscope equipped with an Olympus SC50 camera. The mites were then exposed for three minutes to 3.7 m/s wind containing kairomones derived from either a single plant (Experiment 1), two or three plants (Experiment 2), or no plants (control) (see Table S1 in Supporting Information 1). Then, the number of individuals remaining on the arena was counted. This procedure was repeated five times independently for each generalist and specialist lineages, meaning that for each experimental variant 25 repetitions were performed for specialists and 30 for generalists.

### Statistical analysis

We fitted a beta-binomial generalised linear mixed model (GLMM) using the R package ‘glmmTMB’ (Brooks et al. 2017) to quantitatively assess the considered effects on dispersal rate. Estimated marginal means were used to assess contrasts using the R package ‘emmeans’ (Searle et al. 1980). Analyses were performed using R 4.5 software (R Core Team 2025).

The model testing the contextual effects of olfactory cues included fixed effects for the kairomone (whether it was familiar or not), the current environment in which the mites were placed (familiar/unfamiliar), and host specialisation (specialists or generalists), along with all their interaction terms. It also included a random intercept for the replicates to account for effects that might result from different evolutionary trajectories.

To test for the effect of signal ambiguity, we fitted a model representing the correlation of dispersal rate with the number of kairomones received, separately for each group, which is a combination of a kairomone (using the familiar/unfamiliar split) and host specialisation (specialists or generalists). This was implemented in glmmTMB by specifying the model with the number of kairomones, the above grouping factor and their interaction as fixed effects, and replicates as a random intercept.

The code and data used in the analysis are available on GitHub (https://github.com/popecol/disinfo) and Zenodo (https://doi.org/10.5281/zenodo.16910510).

## RESULTS

### Dispersal in response to kairomones in the context of the current environment and host specialisation

The main effect of kairomones is not significant (p=0.27), which means that there is no general rule in the response to familiar or unfamiliar target plants. When mites are placed in familiar environment, their dispersal rate is lower (0.64 times) compared to an unfamiliar environment (Odds Ratio=0.61, p=0.0005). The overall dispersal rate of the generalist is more than twice that of the specialist (OR=2.37, p<0.0001) (Fig. S3 in Supporting Information 3.1). However, all of the above effects are involved in interactions (Table 1) and their marginal effects should be interpreted with caution, taking into account the possible moderators (see next paragraphs and Supporting Information 2).

**Table 1.** The ANOVA table for a GLMM model examining dispersal rate as a function of the olfactory cue provided by the target environment (herein: Kairomone), the current environment, and host specialisation degree. The p-values less than 0.05 are indicated in bold.

| **Effect** | **$\chi^2$** | **d.f.** | **p-value** |
| --- | --- | --- | --- |
| Kairomone (Control, Familiar or Unfamiliar) | 2.59 | 2 | 0.2735 |
| Current environment (Familiar or Unfamiliar) | 11.60 | 1 | **0.0007** |
| Specialisation (Generalist or Specialist) | 34.32 | 1 | **<0.0001** |
| Kairomone : Current environment | 2.74 | 2 | 0.2540 |
| Kairomone : Specialisation | 14.70 | 2 | **0.0006** |
| Current environment : Specialisation | 19.55 | 1 | **<0.0001** |
| Kairomone : Current environment : Specialisation | 5.06 | 2 | 0.0796 |

Contrary to expectations, the interaction between current and target environment (i.e. the olfactory cue received) is not significant (p=0.25). Irrespective of the type of cues mites received (familiar, unfamiliar or none), they disperse more readily when placed in an unfamiliar current environment (i.e. S and B for specialists; S for generalists), although this difference is not significant for the control, i.e. when no cues are provided (Fig. 1).

**Figure 1.**
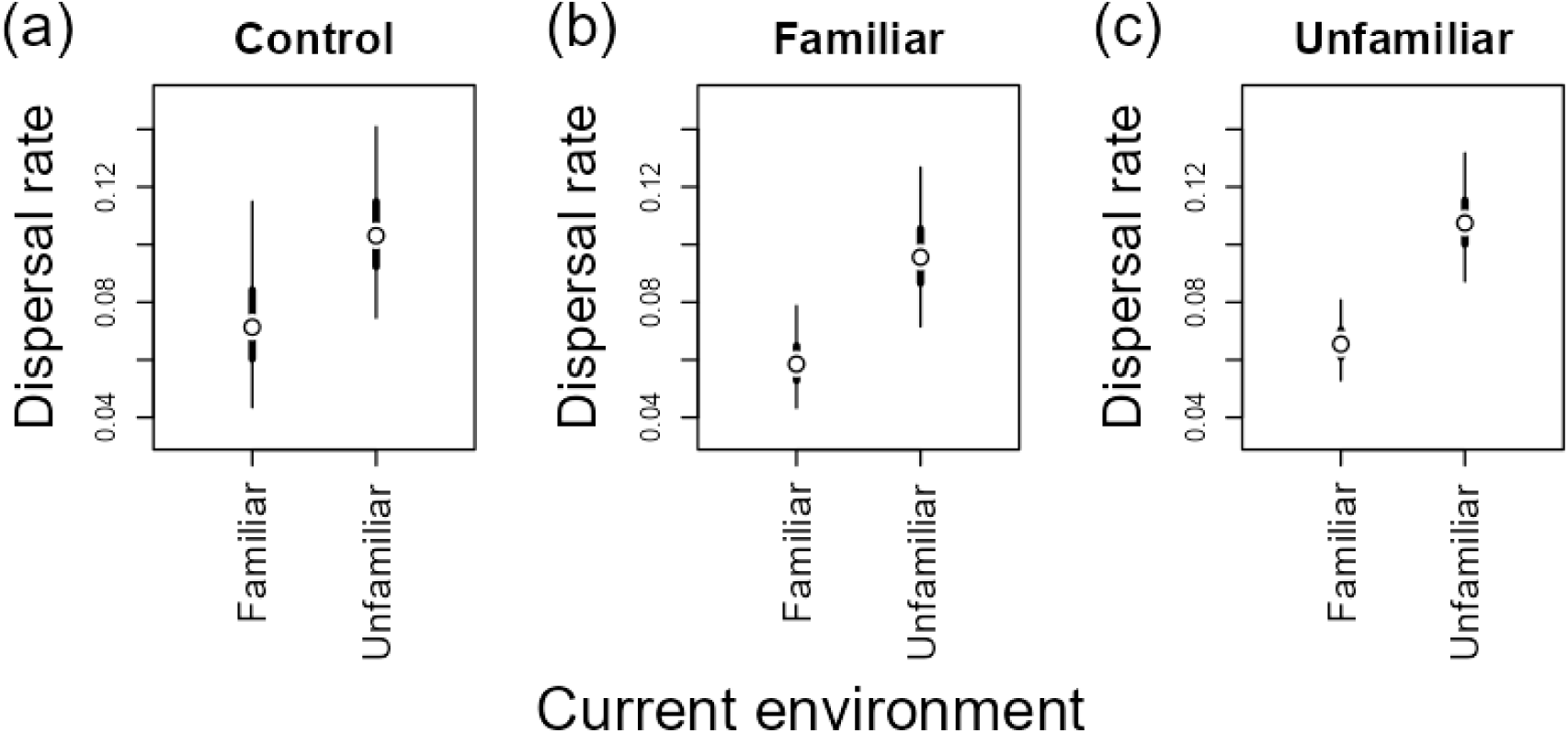
Dispersal rates in relation to the current environment (whether familiar or not) and the type of information received (kairomones) from the target environment. The Control group (a) received no kairomones; the Familiar group (b) received kairomones with which the mites had come into contact during experimental evolution; and the Unfamiliar group (c) received kairomones that they had never encountered before. The points represent the estimated marginal means; the thick and thin lines indicate the 50% and 95% confidence intervals, respectively.

Both the interaction between familiarity of cues and host specialisation and between current environment and host specialisation are significant (p<0.0001 and p=0.0006, respectively). In general, the dispersal rates of generalists are higher than those of specialists, but this difference is much greater when familiar cues are provided. Furthermore, generalists disperse more when the cues are familiar, but the opposite is true for specialists: they disperse more when unfamiliar cues are given. Both generalists and specialists disperse at a rate that does not differ from that of the control (no cues at all), regardless of whether they are familiar or not (Fig. 2). However, an additional analysis at the level of specific host plant species (W, B, S, and O, rather than just the familiar/unfamiliar split) shows differences between specialists and generalists. The specialists’ response to cues from four host species is generally no different from the control, i.e. they respond similarly to cues from familiar and unfamiliar hosts. In contrast, the response of generalists to cues from wheat and barley (familiar hosts) is higher than to the control ((p=0.0286 and p=0.0126, respectively) (see Table S2 and Fig. S4 in Supporting Information 3.2).

**Figure 2.**
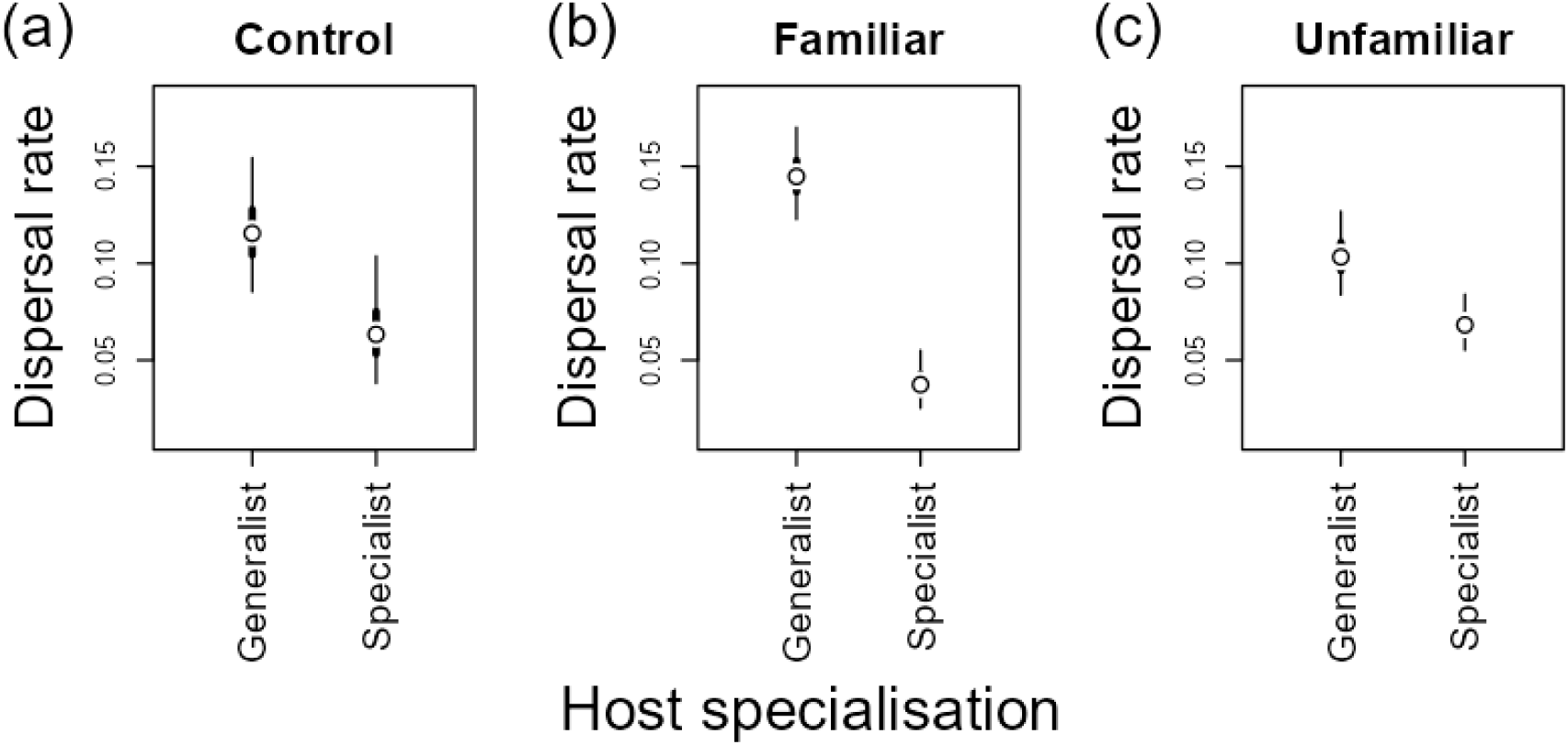
Dispersal rates in relation to host specialisation degree (Generalist vs. Specialist) and the kairomones received from the target environment. The Control group (a) received no kairomones; the Familiar group (b) received kairomones with which the mites had come into contact during experimental evolution; and the Unfamiliar group (c) received kairomones that they had never encountered before. The points represent the estimated marginal means; the thick and thin lines indicate the 50% and 95% confidence intervals, respectively.

Specialists and generalists react differently when placed in a familiar versus an unfamiliar current environment. The dispersal rate of generalists is similar in both environments (p=0.27), whereas specialists disperse more readily in the unfamiliar environment (p<0.0001) (Fig. 3).

**Figure 3.**
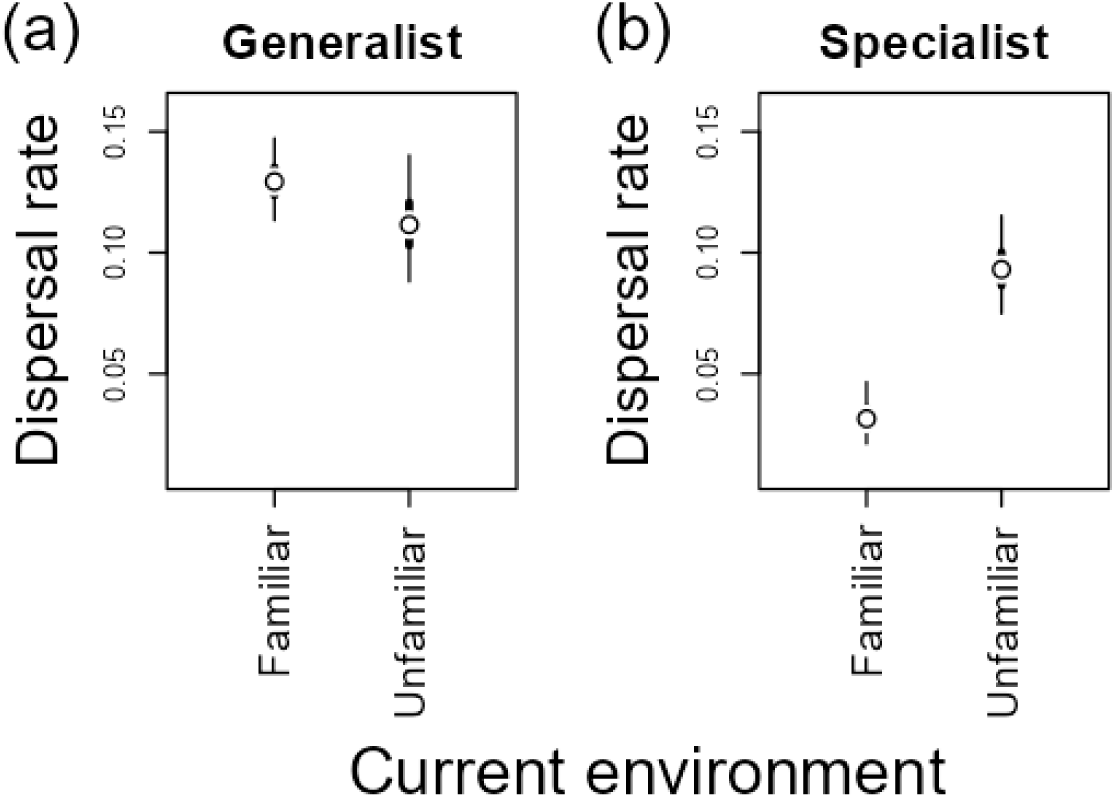
Dispersal rates in relation to the current environment (whether familiar or not) and the host specialisation degree: Generalist (a) vs. Specialist (b). The points represent the estimated marginal means; the thick and thin lines indicate the 50% and 95% confidence intervals, respectively.

The three-way interaction is not significant at the 0.05 level, suggesting that the patterns revealed by the two-way interactions are consistent and not modulated by any of the factors considered.

### Dispersal in response to the number of kairomones in the context of host specialisation

This analysis compares the slopes of dispersal rates against the number of kairomones in the mixture received by different groups. These groups reflect the context of host specialisation, combining the type of olfactory cues provided (familiar or unfamiliar) with the degree of host specialisation (specialists or generalists). Cues are regarded as ‘unfamiliar’ if none of the kairomones in the delivered mixture have been encountered previously. The term ‘familiar’ refers to those instances where at least one of the kairomones has been encountered previously. The only slope that differs from zero is that of the generalists when familiar cues are given. This means that increasing the number of kairomones results in reduced dispersal, but only in generalists and only in response to known kairomones (z=-6.9, p<0.001) (Fig. 4; Table S3 in Supporting Information 3.3).

**Figure 4.**
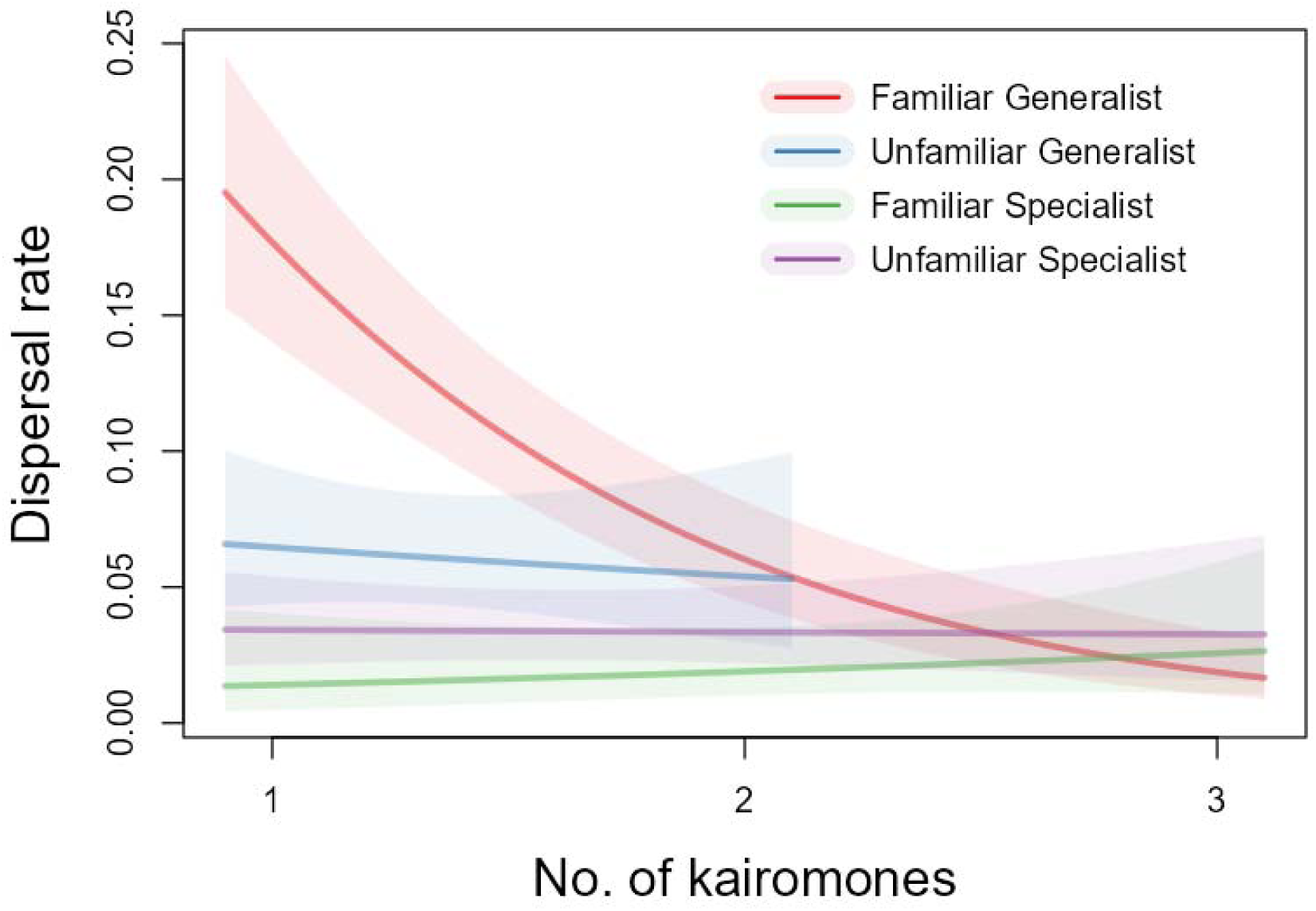
The dispersal rate as a function of the number of kairomones received simultaneously. Lines are fitted for groups representing combinations of cue type (Familiar or Unfamiliar) and host specialisation degree (Generalist or Specialist). Cues are regarded as ‘Unfamiliar’ if none of the kairomones in the delivered mixture have been previously encountered. The term ‘Familiar’ refers to those instances where at least one of the kairomones has been encountered previously. The shaded regions in the figure represent the 95% confidence intervals for the fitted lines.

## DISCUSSION

Passive dispersal relies on external forces, such as wind, water, or other organisms, to enable organisms to move to and colonise new sites. However, even if the subsequent movement is passive, the departure phase, may be at least partially under control and thus actively initiated if influenced by environmental signals. For passively dispersing organisms, kairomones from the environment may function as a reliable source of information to tune dispersal decisions. These chemical substances may indicate a deterioration in the quality of the current habitat, favourable weather conditions, or the presence of more favourable and attractive conditions elsewhere (Bilton et al. 2001; Bonte et al. 2007; Brikett et al. 2004; Bruce et al. 2005; Magalhães et al. 2002; Magalhães et al. 2018).

Our experiments show that the passive dispersal of phytophagous eriophyoid mites is shaped less by the identity of the kairomones per se than by the context in which those cues are encountered and, above all, by the degree of host specialisation. While kairomones alone did not raise departure rates, mites left plants far more readily when they found themselves on an unfamiliar host, and this effect was more pronounced among specialists. Overall, generalists dispersed more than twice as often as specialists. However, their responses diverged sharply in the presence of familiar scents: generalists increased their departures when exposed to any familiar blend, whereas specialists did the opposite, leaving more frequently when the cue came from an unfamiliar plant. Increasing the number of unfamiliar kairomones accentuated the inhibition of generalists dispersal but had little impact on specialists. Together, these patterns suggest that specialists use environmental novelty as a trigger for exploration, whereas generalists rely on multi-cue confirmation of a suitable host before committing to stay. This reveals a fundamental trade-off between the host range and sensitivity to environmental information.

### Information from the current and target environments

Theory predicts that passive dispersers should cling to current familiar habitats due to the high risk of landing on an unsuitable host during random aerial journeys (Bonte et al. 2012; Teller et al. 2015). However, our results only partly bear this out. The wheat-adapted specialist does indeed ‘play it safe’: it remains on wheat, where its fitness is consistently high. This confirms the idea that why not stay put if it is better at home (MacArthur 1972). However, specialists readily take off from unfamiliar barley or smooth brome, where their performance drops (Skoracka et al. 2022). By contrast, generalists show no such home-site loyalty; their baseline departure rate is high for both familiar and unfamiliar plants. This might suggests that generalists do not discriminate between cues from familiar and unfamiliar plants, because receiving cues is costly, either in terms of developing sensory mechanisms and the time needed to perceive them (Dicke 2000). Yet, generalists’ movements are far from indiscriminate, as they do respond to cues from the target environment, as discussed below. Therefore, it is more likely that generalists have a greater chance of finding a suitable environment and face a lower risk of failure than specialists. This is particularly plausible, given that generalists perform well even on plants to which they are not adapted, such as rye and smooth brome (Skoracka et al. 2022). Hence, our experiments suggest that generalists employ a more general bet-hedging strategy, spreading their offspring across the environment, regardless of the information received from the current environment.

In addition to acknowledging the quality of the current environment, identifying signals about the potential future environment could confer an evolutionary advantage (Nichols et al. 2020). Furthermore, the strength and even the direction of the kairomone effect may shift with host specialisation. We expected the wheat-adapted specialists to react most strongly to familiar scents, because making a mistake could cause them to become stranded on a non-host plant. The behavioural activity of phytophagous insects often depends on their perception of blends (Bruce et al. 2005; Bruce & Pickett 2011). However, the overall departure rates of our specialists were lower, actually rising in the presence of unfamiliar cues and falling back to baseline when familiar wheat volatiles were offered. Nevertheless, contrasts at the level of plant species confirmed that the departure rate of specialists remained low, regardless of whether the cue came from wheat, rye, barley, or smooth brome. This suggests that there is a little or no olfactory discrimination (see Fig. S4 in the Supporting Information 3). This pattern implies that it is safest to remain on a known, suitable host plant. Unfamiliar odours may merely signal a change in the landscape, triggering a precautionary take-off – a potentially risky and maladaptive response for a strict specialists.

The generalists showed the opposite tendency. They baseline dispersal in the presence of kairomones was high, and they treated both cue types similarly at the broad ‘familiar/unfamiliar’ level. Yet, the finer plant species-level analysis revealed clear selectivity: cues from wheat and barley (its preferred hosts) boosted departure significantly more that the control, whereas scents from rye or smooth brome did not (see Fig. S4 in the Supporting Information 3). These results mirror those observed in other passive generalists, such as the spider mite, *Tetranychus urticae*, which prefers to move towards the volatiles of its own host plant (Gotoh et al. 1993), and Lepidoptera larvae, which use plant odours to guide their settlement after windborne dispersal (Zalucki et al. 2002). For generalists, the ability to detect a nearby suitable host evidently reduces the risk of aerial exploration. This aligns with the theory that broad-niche organisms can afford, and even benefit, from more frequent movements (Büchi & Vuilleumier 2014; Ravigné et al. 2024; Ward et al. 2001).

We demonstrated that single familiar kairomones stimulate take-off in generalists, whereas blends enriched with unfamiliar cues suppress it. Thus, while specialists treat environmental novelty as a cue to explore, generalists use chemical information to fine-tune risk: they probe when they detect a whiff of a known host, but stay put once multiple matching cues confirm that the current plant is suitable. This decision-making process is consistent with the broader host range of generalists and their ability to maintain fitness even on non-preferred hosts, as demonstrated experimentally (Skoracka et al. 2022).

### Environmental heterogeneity

Natural foragers rarely encounter an isolated host plume; instead, they must extract meaningful information from a chemically crowded backdrop comprising many plant species and herbivores (Beyaert & Hilker 2014; Bruce et al. 2005). In line with field surveys indicating that chemical diversity can dilute herbivore attacks (Salazar et al. 2016), our mixture experiment revealed a significant ‘noise’ effect, albeit only for the generalists. The wheat specialists maintained a uniformly low departure rate regardless of how many plants contributed kairomones. This suggests that they either ignore olfactory background cues or that their decision to leave or stay is governed by cues indicating host quality at the current site. By contrast, the generalists reacted sharply to a single familiar odour; their departure rate was the highest when wheat volatiles were presented alone. That response weakened step-wise as additional, unfamiliar cues were blended in. In other words, the more heterogeneous the signal, the more uncertain the generalists were about whether to emigrate.

One plausible explanation for these results is ratio-specific cue recognition: generalists track the exact proportions of common, shared volatiles in order to distinguish a host plume. Therefore, the addition of non-host odours distorts the diagnostic ratio and erodes the signal (Bruce et al. 2005). Such ratios are inherently difficult to detect amidst a turbulent mosaic of physiologically active compounds. The error would then favour repeated bouts of dispersal until a clean, host-rich plume is found, which indicates a broader matching environment. This explanation aligns with the theory predicting elevated dispersal in broad-niche taxa (Büchi & Vuilleumier 2014; Ravigné et al. 2024; Ward et al. 2001). Alternatively, species-specific markers in the form of unique volatiles for a single host would provide an unambiguous cue. However, evidence for their primacy remains limited (Bruce et al. 2005). Our data therefore support the view that, in noisy landscapes, generalists pay a signal-to-noise penalty, which selects for both rapid departure upon detecting a lone host scent and a readiness to continue searching once that scent is masked by heterospecific odours.

### Degree of host specialisation and response to cues

In our experiments, departure is governed less by the raw presence of kairomones than by the mite’s degree of host specialisation. This is consistent with joint evolution models, in which niche breadth and dispersal form a feedback loop: specialisation is paired with philopatry, whereas a broad diet reduces the costs of “rolling the dice” and therefore selects for high dispersal (Ravigné et al. 2024). Empirically, the wheat-adapted specialists left the plant approximately half as often as their generalist sister lineages (overall OR ≈ 0.42), even when accounting for cue type and current host, confirming the predicted asymmetry. Whether this pattern is universal among phytophagous arthropods remains unclear. Comparative and experimental surveys find that roughly 60% of cases show higher movement in generalists, but there are counter-examples as some specialists – typically those with well-developed movement capacities and perceptual range - disperse more, and many studies report no correlation at all (e.g. Dapporto & Dennis 2013; Dennis et al. 2000; Nieminen 1996; Van Zandt & Mopper 1998). For passively transported taxa, direct tests are rarer still, if exist at all.

The theory also predicts that specialists should rely more heavily on host volatiles in order to compensate for the high cost of making a mistake. Yet, electrophysiological studies often reveal similar receptor breadth in both specialists and generalists (Bruce et al. 2005). Our results echo this paradox: specialists did not respond more strongly to familiar cues, whereas generalists fine-tuned their already high dispersal according to olfactory context. Together, these findings imply that diet breadth shapes the baseline propensity to leave, while chemical information modulates this tendency in a manner that depends on, rather than determines, specialisation. To gain a fuller picture, it will be necessary to track both take-off and settlement in more passive dispersers, and analyse whether the underlying olfactory architecture truly diverges with niche breadth or merely feeds into different decision-making processes.

##### Practical implications for improving pest control

Many herbivorous arthropods that disperse passively, including the wheat curl mite (*Aceria tosichella*), a species we studied here are important agricultural pests. Therefore, understanding the mechanisms underlying their dispersal, particularly those influencing their propensity to depart, is important not only from an ecological and evolutionary perspectives but also for developing crop protection strategies. Such strategies could include engineering or selecting crops for endogenous resistance to pests, or by treating crops with semiochemicals to make them less appealing to pests (Bruce et al. 2005). Specifically, knowledge of which chemical cues discourage and which encourage pest dispersal could potentially help to develop effective, sustainable control and management strategies. For instance, using natural scents derived from plants that pests find unattractive or unfamiliar could restrict their movement towards important crops.

The most important conclusion of our research is that specialists and generalists respond differently to cues from their current and target environments. The experimental conditions, in which evolutionary lineages differing in their degree of host specialisation were obtained, correspond, to some extent, to conditions that might occur in agricultural systems. Specialists with low dispersal rates evolved in the conditions similar to monocultures. Therefore, we can expect such conditions to produce pests that are less likely to disperse to other cereal fields. However, these specialist pests may utilise resources more efficiently than generalists, making them more destructive to crops. Conversely, under conditions conducive to the evolution of generalists (e.g. crop rotation), we can expect pests with high departure rates and thus to be more effective at infesting neighbouring crop fields. It transpires that within a single species, populations can react differently to the level of temporary environmental heterogeneity, which translates into their potential for dispersal and, consequently, their invasive potential. Understanding how agricultural practices influence pests’ propensity to spread can inform the planning of crop protection strategies.

## ACKNOWLEDGMENTS

We are grateful to Alicja Laska-Modzelewska and Anna Przychodzka for their contribution to maintaining the experimental evolution process, to Jarosław Raubic and Grzegorz Zalewski for their invaluable technical support. We also thank the company DANKO Hodowla Roślin in Choryń for providing the seeds of *Triticum aestivum*, *Hordeum vulgare*, and *Secale cereale*, the company CENTNAS in Krotoszyn, and Botanical Garden in Bydgoszcz for the seeds of *Bromus inermis*. The study was supported by the National Science Centre (NSC), Poland grant no. 2019/35/N/NZ8/03377 to KZ. AS was involved in this work while supported by NSC grant no. 2021/41/B/NZ8/01703.

All authors declare no conflict of interest.

## Statement of authorship

KZ conceptualized and designed the study with LK, AS, EP, ML assistance; KZ ran experiments and collected data; LK performed statistical analyses; LK, AS, DB, EP, ML and KZ critically interpreted the results; AS, LK and DB wrote the manuscript with assistance of EP, ML, KZ; All authors contributed substantially to revisions, read and approved the final version of the manuscript.

## Data accessibility

The data that support the findings and the code used to produce the results of this study are openly available on GitHub: https://github.com/popecol/disinfo and Zenodo: https://doi.org/10.5281/zenodo.16910510

