## Supporting Information for "Is passive dispersal informed? - Experimental evidence for decision-making in phytophagous arthropods"

### Supporting Information 1 – Supplementary Methods

**Table S1**. Experimental variants made to test the dispersal rate of the wheat curl mite (WCM) in the response to cues from the current and target environments consisting of plant species: B=barley, O=oats S=smooth brome, W=wheat, and 0=no cues.

| Experiment | Current environment | Target environment | Remarks |
|---|---|---|---|
| 1 | W | 0 | Control |
| 1 | S | 0 | Control |
| 1 | B | 0 | Control |
| 1 & 2 | W | W | 1 kairomone |
| 1 & 2 | W | B | 1 kairomone |
| 1 & 2 | W | S | 1 kairomone |
| 1 & 2 | W | O | 1 kairomone |
| 1 | B | W | 1 kairomone |
| 1 | B | B | 1 kairomone |
| 1 | B | S | 1 kairomone |
| 1 | B | O | 1 kairomone |
| 1 | S | W | 1 kairomone |
| 1 | S | B | 1 kairomone |
| 1 | S | S | 1 kairomone |
| 1 | S | O | 1 kairomone |
| 2 | W | W-S | 2 kairomones |
| 2 | W | S-O | 2 kairomones |
| 2 | W | B-S-O | 3 kairomones |
| 2 | W | W-S-O | 3 kairomones |

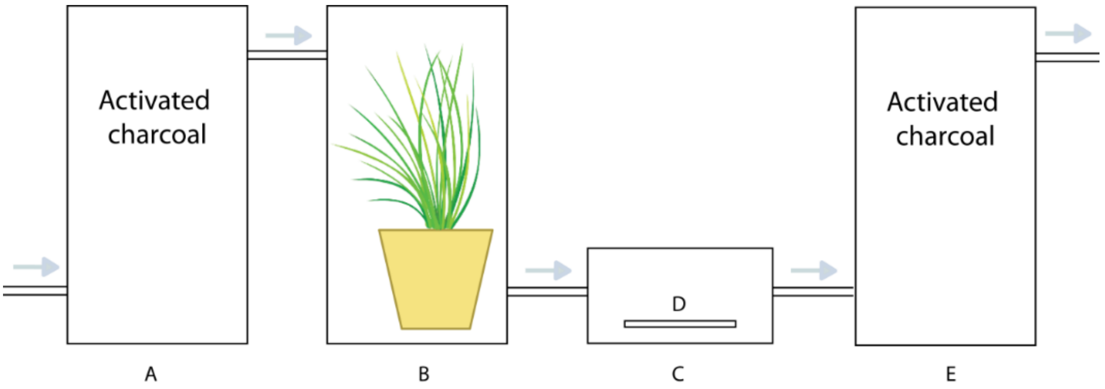


**Fig. S1**. Schematic diagram of the olfactometer. A, E – chamber filled with activated charcoal; B – chamber with a plant being the source of kairomones; C – chamber in which tested wheat curl mite (WCM) individuals were placed; D – plant fragment with mite individuals placed on agar blocks: experimental arena – current environment.

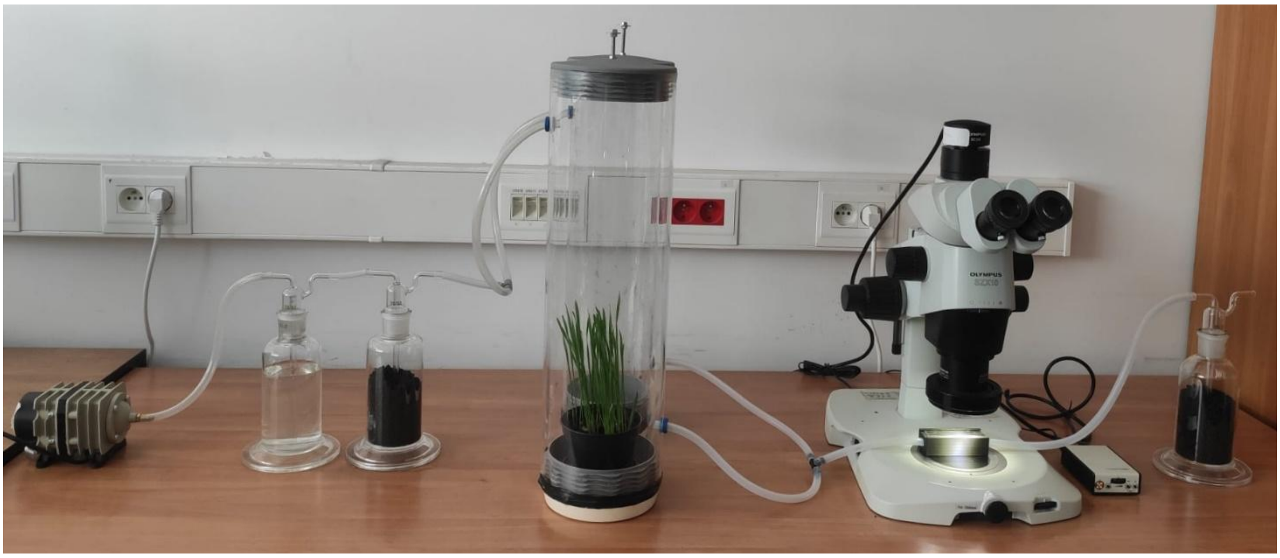

**Fig. S2**. The equipment for testing dispersal responses to kairomones (olfactometer).

### Supporting Information 2 – Estimated marginal means contrasts

#### 1. Marginal effects

##### 1.1. Kairomones (cues)

```
contrast               odds.ratio    SE  df null z.ratio p.value
 control / familiar         1.159 0.228 Inf    1   0.748  0.7347
 control / unfamiliar       1.023 0.184 Inf    1   0.127  0.9911
 familiar / unfamiliar      0.883 0.123 Inf    1  -0.892  0.6451

Results are averaged over the levels of: env, host_spec
P value adjustment: tukey method for comparing a family of 3 estimates
Tests are performed on the log odds ratio scale
```

##### 1.2. Current environment

```
contrast               odds.ratio     SE  df null z.ratio p.value
 familiar / unfamiliar      0.612 0.0869 Inf    1  -3.462  0.0005

Results are averaged over the levels of: cue, host_spec
Tests are performed on the log odds ratio scale
```

##### 1.3. Specialisation

```
contrast odds.ratio    SE  df null z.ratio p.value
 G / S         2.37 0.337 Inf    1   6.056  <.0001

Results are averaged over the levels of: cue, env
Tests are performed on the log odds ratio scale
```

#### 2. Conditional effects: 2-way interactions

##### 2.1. Kairomones : Current environment

```
cue = control:
 contrast               odds.ratio     SE  df null z.ratio p.value
 familiar / unfamiliar      0.669 0.2150 Inf    1  -1.251  0.2110

cue = familiar:
 contrast               odds.ratio     SE  df null z.ratio p.value
 familiar / unfamiliar      0.588 0.1340 Inf    1  -2.338  0.0194

cue = unfamiliar:
 contrast               odds.ratio     SE  df null z.ratio p.value
 familiar / unfamiliar      0.582 0.0944 Inf    1  -3.339  0.0008

Results are averaged over the levels of: host_spec
Tests are performed on the log odds ratio scale
```

##### 2.2. Kairomones : Specialisation

```
cue = control:
 contrast odds.ratio    SE  df null z.ratio p.value
 G / S          1.93 0.622 Inf    1   2.044  0.0409

cue = familiar:
 contrast odds.ratio    SE  df null z.ratio p.value
 G / S          4.36 0.993 Inf    1   6.473  <.0001

cue = unfamiliar:
 contrast odds.ratio    SE  df null z.ratio p.value
 G / S          1.57 0.255 Inf    1   2.794  0.0052

Results are averaged over the levels of: env
Tests are performed on the log odds ratio scale
```

##### 2.3. Current environment : Specialisation

```
host_spec = G:
 contrast              odds.ratio     SE  df null z.ratio p.value
 familiar / unfamiliar      1.183 0.1800 Inf    1   1.108  0.2677

host_spec = S:
 contrast              odds.ratio     SE  df null z.ratio p.value
 familiar / unfamiliar      0.316 0.0759 Inf    1  -4.798  <.0001

Results are averaged over the levels of: cue
Tests are performed on the log odds ratio scale
```

#### 3. Conditional effects: 3-way interactions

```
cue = control, host_spec = G:
 contrast              odds.ratio     SE  df null z.ratio p.value
 familiar / unfamiliar      2.170 0.7390 Inf    1   2.276  0.0228

cue = familiar, host_spec = G:
 contrast              odds.ratio     SE  df null z.ratio p.value
 familiar / unfamiliar      0.797 0.1540 Inf    1  -1.172  0.2413

cue = unfamiliar, host_spec = G:
 contrast              odds.ratio     SE  df null z.ratio p.value
 familiar / unfamiliar      0.958 0.2240 Inf    1  -0.184  0.8539

cue = control, host_spec = S:
 contrast              odds.ratio     SE  df null z.ratio p.value
 familiar / unfamiliar      0.206 0.1130 Inf    1  -2.890  0.0039

cue = familiar, host_spec = S:
 contrast              odds.ratio     SE  df null z.ratio p.value
 familiar / unfamiliar      0.434 0.1780 Inf    1  -2.032  0.0421

cue = unfamiliar, host_spec = S:
 contrast              odds.ratio     SE  df null z.ratio p.value
 familiar / unfamiliar      0.353 0.0794 Inf    1  -4.632  <.0001

Tests are performed on the log odds ratio scale
```

### Supporting Information 3 – Supplementary Results

#### 1. The main effect of kairomones

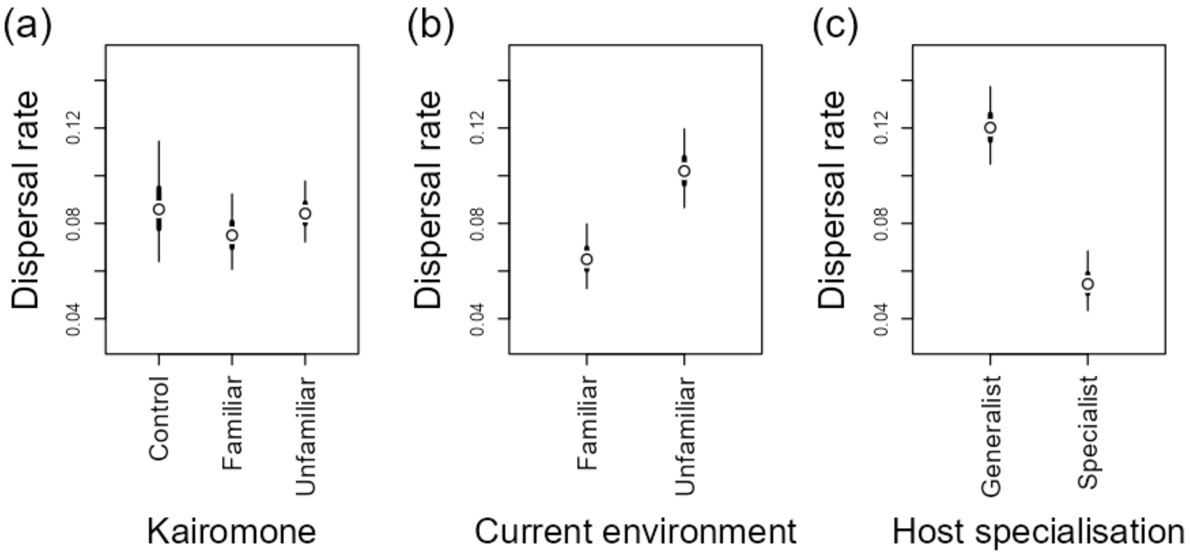


**Fig. S3**. Main effects of kairomones (a), current environment (b) and host specialisation (c) on dispersal rates. The points represent the estimated marginal means; the thick and thin lines indicate the 50% and 95% confidence intervals, respectively.

#### 2. The analysis performed on specific plant species

This analysis generally confirmed the results obtained from the analysis grouping familiar versus unfamiliar plant species (Table S2). Furthermore, it clarified the interpretation of generalists and specialists responses to cues from the target environment (see Fig. 2 in the main manuscript). The dispersal rates of generalists in the presence of cues from familiar hosts (wheat and barley) are significantly higher than in the control (p=0.0286 and p=0.0126, respectively) as well as in the presence of cues from smooth brome (p<0.001 for both contrasts). This suggests that generalists recognise cues from familiar hosts. However, generalists dispersal rates in the presence of cues from unfamiliar plants (smooth brome and rye) do not differ significantly from the control (p=0.1672 and p=0.9874, respectively). The dispersal rates of specialists are generally low and do not differ significantly in the presence of kairomones from all plant species (wheat and unfamiliar ones) and the control (p>0.1) (Fig. S4).

**Table S2**. The ANOVA table for a GLMM model examining dispersal rate as a function of olfactory cue provided (a kairomone emitted by a specific host plant species), current environment (current host plant species) and host specialisation (specialists or generalists).

| **Effect** | $\chi^2$ | **d.f.** | **p-value** |
|---|---|---|---|
| Kairomone (4 plant species + control) | 16.7 | 4 | **0.0022** |
| Environment (3 plant species) | 61.9 | 2 | **<0.0001** |
| Specialisation (generalists or specialists) | 31.4 | 1 | **<0.0001** |
| Kairomone : Environment | 10.2 | 8 | 0.2561 |
| Kairomone : Specialisation | 30.2 | 4 | **<0.0001** |
| Environment : Specialisation | 8.6 | 2 | **0.0137** |
| Kairomone : Environment : Specialisation | 10.2 | 8 | 0.2499 |

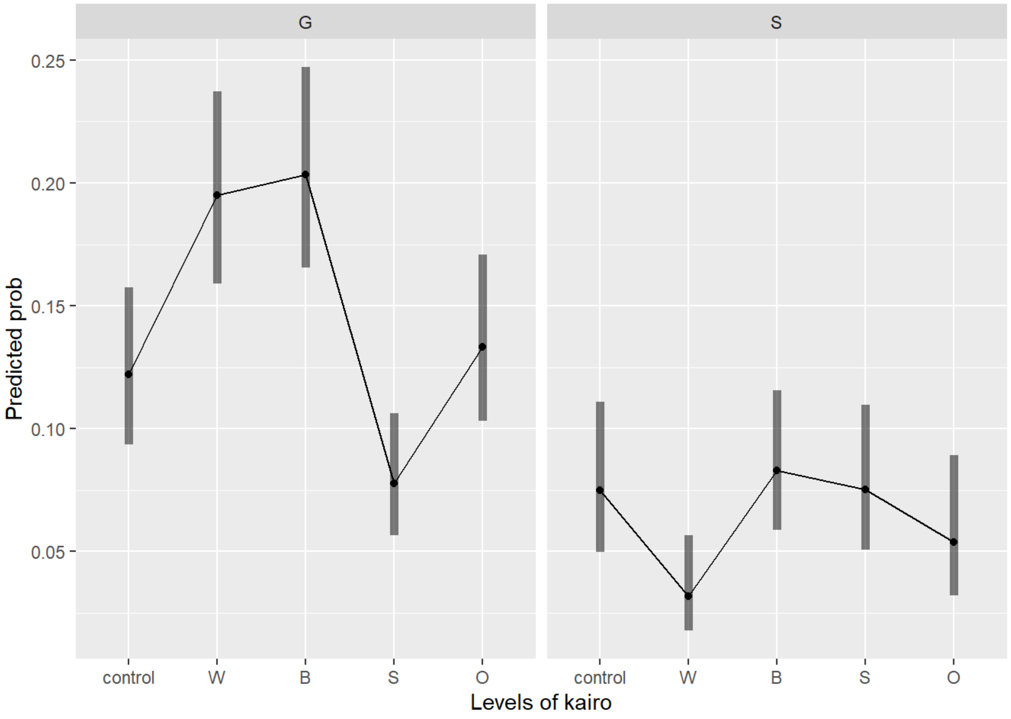


**Fig. S4**. 2-way interactions: dispersal rates in relation to cues received (panels: control=none, W=wheat, B=barley, S= smooth brome, O=oat) and host specialisation (G=generalists, S=specialists).

#### 3. Dispersal in response to the number of kairomones

Table S3. Parameters of the GLMM model relating the number of kairomones to the dispersal rate. Lines are fitted separately for each combination of kairomone familiarity and host specialisation.

| | **Estimate** | **SE** | **z-value** | **p-value** |
|---|---|---|---|---|
| **Intercepts** | | | | |
| Generalist-Familiar host | -0.33 | 0.27 | -1.21 | 0.2250 |
| Generalist-Unfamiliar host | -2.48 | 0.52 | -4.80 | **<0.0001** |
| Specialist-Familiar host | -4.56 | 0.90 | -5.09 | **<0.0001** |
| Specialist-Unfamiliar host | -3.31 | 0.42 | -7.82 | **<0.0001** |
| **Slopes (against number of kairomones)** | | | | |
| Generalist-Familiar host | -1.21 | 0.17 | -6.92 | **<0.0001** |
| Generalist-Unfamiliar host | -0.19 | 0.37 | -0.52 | 0.6050 |
| Specialist-Familiar host | 0.31 | 0.38 | 0.80 | 0.4210 |
| Specialist-Unfamiliar host | -0.02 | 0.23 | -0.11 | 0.9160 |
